# Characterization of a virulent bacteriophage consortium targeting Enterobacteriaceae from inflamed preterm gut mucosa

**DOI:** 10.1101/2025.01.29.635513

**Authors:** Malene Roed Spiegelhauer, Simone Margaard Offersen, Lukasz Krych, Dennis Sandris Nielsen, Michela Gambino, Anders Brunse, Duc Ninh Nguyen

## Abstract

Preterm infants have a high risk of intestinal inflammation which can progress to necrotizing enterocolitis (NEC). The gut microbial colonization commencing at birth is essential for proper intestinal maturation, but this process is often disrupted in preterm infants, leading to dysbiosis and increased risk of developing NEC. Bacteriophages (phages), viruses that specifically infect bacteria, are an important constituent of the gut microbiome and protects the gut epithelium against invading bacteria. This study aimed to isolate and characterize phages for use as a preventive measure against NEC-associated bacteria. We initially cultured *Enterobacteriaceae* from ileal mucosa of preterm piglets that exhibited severe NEC-like pathology. We then screened 23 donor fecal samples for inhibition of bacterial growth and isolated a collection of unique phages to use further. The phages were characterized by whole genome sequencing, host receptor binding determination, and immune cell activation *in vitro*. The final phage collection consisted of ten virulent phages within five genera, representing myovirus, podovirus and siphovirus morphologies. All phages in the collection induced expression of both pro-and anti-inflammatory genes in co-culture with macrophage-like THP-1 cells, but to different extents than *Escherichia coli*. Ultimately, we selected one phage for a high-dose oral administration to newborn piglets and assessed its infectivity and presence in different gut segments. This intervention did not result in any direct side effects, while both infective phages and signatures of phage DNA were detected in the intestinal content and mucosa. Having characterized a set of rationally selected virulent phages, we support the advancement of phage therapy as a potential protection against NEC.

## Introduction

Preterm birth, defined as birth before 37 weeks’ gestation and affecting 15 million infants each year, is the leading cause of neonatal mortality and long-term morbidities [1]. The preterm intestine is predisposed to hyper-inflammatory mucosal responses and development of necrotizing enterocolitis (NEC), a severe intestinal disease with high mortality [2]. Establishment of a diverse gut microbiome is crucial for intestinal maturation and improved barrier function [3]. This assembly appears to be impeded in preterm infants and leads to dysbiosis characterized by dominance of facultative anaerobic bacteria and a delay in the natural transition to obligate anaerobes [4]. The preterm gut microbiome is additionally characterized by low abundance of e.g. *Bifidobacterium* members which are normally found in high numbers in term infants, and increase in *Enterobacteriaceae*, a family of Gram-negative bacteria that includes several pathogens [5]. Specifically, infants developing NEC have a higher abundance of bacteria belonging to *Enterobacteriaceae* such as *Escherichia coli, Klebsiella pneumoniae* or *Enterobacter cloacae* prior to diagnosis, suggesting that this dysbiosis might contribute to the disease development [6], [7].

The gut contains a diverse array of bacteriophages (phages), viruses that infect bacteria in a highly specific manner. Phages initiate infection by binding to a specific structure on the bacterial host surface and transferring its genome into the bacterial cell. In virulent phages, this is immediately followed by the production of new phage particles which are ultimately released through bacterial lysis [8]. In addition to this, temperate phages can instead integrate their genomic material into the bacterial genome, where they remain dormant until induced [8]. In parallel with gut bacterial colonization, a phage community establish during the first weeks of life, quickly reaching a density of up to 10^9^ phage particles per gram of feces [9], [10], [11]. Mechanistically, phages can accumulate in the intestinal mucus layer, where they protect the underlying epithelium against invasive bacteria [12]. The roles of gut phages and phage-bacteria interactions in the newborn gut, as well as their contributions to infant health, are largely unexplored.

A recent observational study revealed crucial fluctuations in the preterm infant gut virome prior to the onset of NEC, characterized by altered composition, reduced interpersonal variation and the presence of viral species associated with the development of NEC [13]. Consequently, modulation of the preterm gut virome may hold potential for preventing NEC. We have previously demonstrated a protective effect of transferring sterile donor fecal filtrate (FFT), containing a diverse phageome, to NEC-susceptible preterm piglets [14]. Importantly, microbiome analysis revealed an increased viral diversity and reduced relative abundance of bacteria belonging to *Enterobacteriaceae* in ileal mucosa of piglets receiving fecal filtrate, supporting the notion that the protective mechanism may be mediated by exogenous phages located in the gut mucosa.

This study aimed to obtain a collection of individual phages with potential to use as a simplified alternative to FFT. We hypothesized that phages specifically infecting bacteria from NEC-like intestinal lesions could be isolated from feces of healthy individuals, and that they would populate the gut mucosa in recipients. To test this hypothesis, we screened fecal filtrates from healthy pigs against a library of *Enterobacteriaceae* cultured from mucosa of active NEC-like lesions of preterm piglets. We successfully isolated a collection of ten phages which were characterized through genomic sequencing and analysis, electron microscopy and host receptor identification. Additionally, we measured their immune stimulating potential *in vitro* and finally trailed the gut distribution of a single phage following oral administration in newborn piglets.

## Materials and Methods

### Mucosa collection and bacterial isolation from mucosa of NEC-like lesions

Fresh ileal sections of NEC-like lesions (displaying extraluminal gas accumulations and necrosis) were collected from 11 formula-fed preterm piglets [15] and flushed with sterile water. The mucosa was scraped off and suspended in 2 ml 10% glycerol, snap-frozen and stored at -80 °C until use. Serial dilutions of the collected mucosa were prepared in 0.9% sodium chloride and cultured aerobically on MacConkey agar (Merck) at 37 °C for 16 hours. The bacterial load was determined as the number of colony-forming units (CFU) per ml suspended mucosa, with a detection limit of 10 CFU/ml. Selected colonies of each morphology were isolated and identified by Matrix-Assisted Laser Desorption Ionised Time Of Flight Mass Spectrometry (MALDI-TOF MS, Bruker).

### Pulsed-Field Gel Electrophoresis

The bacterial isolates were grouped by pulse-field gel electrophoresis (PFGE) following an adapted protocol from PulseNet and Centers for Disease Control [16]. In brief, bacterial cultures were cast in 0.1% agar plugs and incubated at 56 °C with buffer containing 50 mM Tris-HCl, 50 mM EDTA, 1% Sarcosyl (N-Lauroylsarcosine, Sigma-Aldrich), and 25 µl Proteinase K (Invitrogen, Thermo Fisher). The plugs were incubated with the restriction enzyme XbaI (New England Biolabs) in Cutsmart Buffer (New England Biolabs) for 2 hours at 37 °C and stored in Tris-HCl:EDTA 10:1 buffer at 4 °C. As a marker of size, *Salmonella* serotype *Branderup* ladder was digested with XbaI (New England Biolabs) and prepared similar to the samples. The plugs were then embedded in a gel with 1.5% agarose (SeaKem Gold agarose, Lonza), and PFGE was run with the following setup: 0.5X TBE buffer and thiourea, initial switch time 2.2 seconds; end switch time 54.4 seconds; 18 hours runtime; 6V; included angle 120 (CHEF-DR III Pulsed Field Electrophoresis System, Bio-Rad Laboratories). The gel was stained with Ethidium bromide and bands were visualized with the Gel Doc XR+ system (BioRad Laboratories) and Gel Compar II (version 4.6, Applied Maths BVBA).

The gel bands were compared, and different patterns indicated unique bacterial profiles (Supplementary Table S1, Supplementary Figure S1).

### Bacterial isolate whole genome sequencing

Bacterial DNA was extracted with the DNeasy Blood and Tissue kit (Qiagen) and sequenced by Illumina Miseq with 2x300bp paired-end sequencing, and Flex DNA library preparation kit with UD indexes set A for 96 samples. The reads were assembled from FASTq-files by the Comprehensive Genome Analysis tool in the online platform Bacterial and Viral Bioinformatics Resource Center (BV-BRC, www.bv-brc.org) with Unicycler v0.4.8. The species identification was confirmed by aligning the sequenced genomes with the reference genomes of *Escherichia coli* str K-12 substrain MG1655 (NC_000913.3), *Escherichia coli* O157:H7 strain (NC_002695.2) and *E. hormaechi* (*Enterobacter hormaechei subsp. steigerwaltii* (CP100388.1) comparing the Average Nucleotide Identity (ANI) [17]. The isolates were further characterized by prediction of multi-locus sequence typing (MLST) (MLST2.0 [18], [19]), serotypes (SerotypeFinder [20]), virulence genes (VirulenceFinder version 2.0.5 [19], [21], [22]) and antibiotic resistance (ResFinder version 4.5.0 [19], [23]).

### Donor fecal filtrate preparation

Twenty-three porcine fecal filtrates (Supplementary Table S2) were collected as natural phage-rich sources. *Filtrate 1* was prepared in a prior study from pooled colon content of five healthy 10-day-old piglets [14]. The pooled colon content was suspended in sterile 20% glycerol and further diluted 1:10 in sterile SM buffer (200 mM NaCl, 10 mM MgSO4, 50 mM Tris-HCl) with 10% glycerol. The suspension was centrifuged at 5000 × *g* for 30 minutes at 4 °C, and the supernatant was filtered with pore size 0.45 µm (MiniSart Syringe Filter, Sartorius) to obtain the final filtrate, which was stored at -80 °C until use. *Filtrates 2 to18* were prepared from colon contents of healthy piglets between 8 and 12 days old, reared at different conventional and organic Danish farms. The colon contents were diluted 1:20 in sterile SM buffer and homogenized by stomaching, followed by two rounds of centrifugation at 11.000 × *g* for 30 minutes. The supernatant was filtered twice with pore size 0.45 µm (MiniSart Syringe Filter, Sartorius), and stored at -80 °C until use.

*Filtrates 19-23* were prepared in a prior study [24]. In brief, feces from healthy piglets or sows were diluted 1:10 in sterile SM buffer supplemented with 0.01% gelatine. The contents were stomached and centrifuged at 18.000 × *g* for 10 minutes, and the supernatant was passed through a 0.45 µm filter (MiniSart Syringe Filter, Sartorius) twice. Thesefiltrates were stored at 4 °C until use.

### In vitro filtrate screening

Donor fecal filtrates were screened in triplicate wells for bacteriolytic activity against the bacterial isolates. Individual filtrates were mixed 1:1 with a bacterial inoculum (cultured for 2-3 hours) in MacConkey Broth (Merck) on a 96-well plate and co-incubated for 18 hours at 37 °C with continuous shaking and measure of optical density (OD) at 655 nm every 20 minutes (Gen5 Microplate Reader, Imager Software Version 1.10.8, BioTek Instruments, Agilent technologies). The bacterial growth was compared to an inoculum without filtrate. Wells showing growth inhibition in at least one replicate were collected and stored at 4 °C for phage isolation.

### Phage isolation

Phages were isolated from the collected wells using the double agar overlay method [25]. In brief, the well contents were mixed 1:1 or 2:1 with the corresponding bacterial inoculum (cultured for 2-3 hours) and 3.3 ml top agar (BD DIFCO LB Broth, Lennox and 0.6% Oxoid Agar Bacteriological, Thermo Scientific, Lennox), distributed on top of LB agar (BD DIFCO LB agar, Lennox) and incubated at 37 °C. Zones of bacterial lysis (plaques) with unique morphology were picked with a sterile pipette tip, transferred to SM buffer, and incubated for 30 minutes at 37 °C, 450 rpm, to release phages from the agar. If no visible plaques were formed, the plate contents were collected in SM buffer for at least 4 hours at 4 °C, 150 rpm, passed through a 0.45 µm filter (MiniSart Syringe Filter, Sartorius), and incubated with a mix of 10% polyethylene glycol and 0.5 M NaCl at 4°C. The solution was centrifuged at 10.000 × *g* for 1 hour at 4 °C, the pellet was resuspended in 0.1x volume SM buffer and plated again with the double agar overlay method. Single plaques were picked for at least three repetitions of the double agar overlay.

### Phage propagation

The phages were propagated by the double agar overlay method using 100µl phage suspension with a titer between 10^4^-10^5^ plaque-forming units (PFU) per ml and 100 µl host bacterial inoculum (cultured for 2-3 hours), mixed with 3.3 ml top agar distributed on top of LB agar and incubated at 37 °C. To collect phages, 5 ml of SM buffer was added to the plate and incubated at 4 °C, 100 rpm. The liquid was collected, centrifuged at 7000 rpm for 5 minutes and the supernatant was passed through a 0.45 µm filter (MiniSart Syringe Filter, Sartorius). The phage titer was determined by spotting 10-fold dilutions in triplicates of 10 µl on a bacterial lawn prepared with the double agar overlay method, and with a detection limit of 33 PFU/ml.

### Phage DNA extraction

DNA was extracted using a modified phenol-chloroform protocol with ethanol precipitation as described previously [26]. High-titer phage propagations (>10^8^ PFU/ml) were passed through a 0.2 µm filter (MiniSart Syringe Filter, Sartorius) and incubated with 10 µg/ml RNase A (Thermo Scientific) and 20 µg/ml DNase I (Thermo Scientific) for 40 minutes at 37 °C , 500 rpm in a thermoshaker (Eppendorf), followed by incubation at 56 °C with added 20 mM EDTA (pH 8.0) and 50 µg/ml Proteinase K (Invitrogen) to inactivate the nucleases. The extraction consisted of phenol (Sigma-Aldrich), phenol-chloroform-isoamyl alcohol (25:24:1, Sigma-Aldrich), and two rounds of chloroform-isoamyl alcohol (24:1, Sigma-Aldrich) treatment. The DNA was precipitated at -20 °C in a mix of 3 M sodium acetate, 20 mg/ml glycogen (Thermo Scientific), and 99% ethanol, centrifuged at 11.000 rpm at 4 °C and washed three times with 70 % ethanol. After the final wash and centrifugation, the DNA pellets were dried at 37 °C, resuspended in DNAse-free water, and stored at -20 °C. The presence of DNA was confirmed by gel electrophoresis on a 1% agarose gel, and visualized with the Gel Doc XR+ system (Biorad Laboratories) using the program Gel Compar II (version 4.6, Applied Maths BVBA). The DNA quality and quantity were assessed by the Nanodrop and Qubit dsDNA High Sensitivity Assay (Invitrogen) (DS-11 FX+ spectrophotometer/fluorometer, DeNovix Inc.). Samples with low quality were processed with a DNA cleanup kit (Genomic DNA Clean & Concentrator-25, Zymo research) following the manufacturer’s protocol.

### Phage genome sequencing

Seven phages were sequenced by SeqCenter (Pittsburgh, PA) using Illumina sequencing following their pipeline. The genomes were assembled from FASTq-files to FASTA sequences using the Comprehensive Genome Analysis tool (BV-BRC, www.bv-brc.org). Fourteen phages were sequenced at the Department of Food Science at University of Copenhagen, using Nanopore technology. In brief, DNA concentration was normalized to 20 ng/μl (200 ng total) and subjected to library preparation using the Rapid Barcoding Sequencing protocol following the manufacturer’s instructions (SQK-RBK114.24, version RBK_9176_v114_revG_27Nov2022, ONT, Oxford, UK). The DNA concentration was normalized to 50 ng/μl (400 ng total) and loaded on a single R10.4.1 flow cell. The sequencing was performed on the GridION sequencing platform (MinKNOW 21.05.20) and basecalled with Dorado (version 0.5.0, ONT, Oxford, UK). Quality was checked with NanoPlot (version 1.43.0), and genomes were assembled to FASTA sequences with Flye (version 2.9.5-b1801).

### Comparative analyses of sequenced phage genomes

The phage genomes were aligned with BLAST (NCBI) to identify phages with high sequence similarity. The genomic similarities between the isolated phages or between the isolated phages and representative phages for each genus were assessed with the web tool VIRIDIC [27], and tthe intergenomic distances was determined with the web-based tool VICTOR [28] using Herpes Simplex virus type 1 as outgroup. Branch support was inferred by FASTME and SPR postprocessing [29], and phylogenetic trees were visualized from the D0 (nucleotide) algorithm using iTOL with rooted midpoint of tree [30]. The phage genomes were annotated via the Genome Annotation tool (BV-BRC, www.bv-brc.org), with the PHANOTATE recipe, and the taxonomy of the genus reference phage with highest similarity. The genomes were checked for presence of integrase (from the annotated genome), for known virulence genes (VirulenceFinder version 2.0.5 [19], [21], [22]) and antibiotic resistance (ResFinder version 4.5.0 [19], [23]).

### Transmission Electron Microscopy (TEM)

High-titer phage suspensions (>10^8^ PFU/ml) were prepared for electron microscopy following a previously published protocol [31]. In brief, the phages were pelleted by centrifugation at 18.000 x *g* for 60 minutes, resuspended in 0.1x volume of 0.1 M ammonium acetate and stored at 4 °C until analysis. The phages were transferred to 200-mesh copper-coated glow discharged grids (15 mA for 30 seconds, Leica coater ACE 200) and coated with 2% uranyl acetate as negative stain. The phages were visualized at 165.000x-250.000x magnification with the electron microscope PHILIPS FEI CM100, and images were taken with a Veleta side-mounted camera (Olympus) using the iTEM imaging platform (Olympus). The phage dimensions were assessed from the TEM images by five measurements per phage with the ImageJ software [32].

### Receptor binding

The *E. coli* MG1655 wildtype (WT) and its surface protein deletion mutants were used to investigate the receptor binding *(Unpublished, manuscript in preparation)*. Bacterial lawns were prepared for each strain, and the collected phages were individually spotted in serial concentrations between 10^2^ to 10^8^ PFU/ml in triplicates of 10 µl. Phage infectivity was evaluated as the observed phage titer relative to the titer on *E. coli* MG1655 WT, and reduced infectivity was defined as minimum a 1-log reduction of phage titer. Phages unable to infect the *E. coli* MG1655 WT strain were not spotted on the *E. coli* MG1655 mutant strains.

### Endotoxin removal for in vitro stimulation

The ten phages were propagated and passed through an endotoxin removal column according to the manufacturer’s instructions (Pierce™ High-Capacity Endotoxin Removal Spin Columns, Thermo Scientific), and endotoxin levels were confirmed before and after column treatment (Pierce™ Chromogenic Endotoxin Quant Kit, Thermo Scientific).

### Cell culture conditions and in vitro stimulation with phages

THP-1 monocytes were cultured following a previous protocol [33] in growth medium consisting of RPMI 1640 GLUTAMAX (Gibco, Thermo Scientific) supplemented with 10% heat-inactivated fetal bovine serum (FBS, Gibco, Thermo Scientific) and 25 µg/ml gentamicin (Gibco, Thermo Fisher) and incubated at 37 °C, 5% carbon dioxide. The monocytes were differentiated into macrophages by seeding 2 x 10^6^ cells per well in a 12-well plate in growth medium supplemented with 5 ng/µl phorbol 12-myristate 13-acetate (Thermo Fisher Scientific) for 48 hours. Before co-culture, the medium was replaced with RPMI 1640 GLUTAMAX supplemented with only 10% heat-inactivated FBS, and the cells were incubated for an additional hour. The co-cultures were performed in biological triplicates using different cell passages. Phages were added to wells individually in a titer of 10-100 PFU per cell, and as a reference of response against bacteria, the *E. coli* ST-2064 was prepared and added in a multiplicity of infection at 10 bacteria per cell following a previous protocol [33]. Cells co-cultured with sterile SM buffer were used as control for unchallenged cells. The co-culture plates were centrifuged at 300 rpm for 5 minutes and incubated for 20 hours at 37 °C, 5% carbon dioxide.

### RNA extraction, reverse transcription and qPCR

RNA was extracted from cell cultures with the RNeasy Mini Kit (QIAGEN) according to the manufacturer’s instructions and including DNase treatment (RNase-free DNase, QIAGEN). RNA quantity and quality were measured on NanoDrop ND-1000 (Thermo Fisher), and 10 ng RNA was reverse transcribed to cDNA (High-Capacity cDNA Reverse Transcription kit, Applied Biosystems). Duplicate reactions of quantitative PCR (qPCR) were prepared in 10 µl volumes with 1 x SYBR Green Master (LightCycler 480 SYBR Green Master, Roche), 0.4 µM forward primer and 0.4 µM reverse primer (TAG Copenhagen). The cycling conditions were: 15 minutes at 95 °C, 45 cycles of (15 seconds at 94 °C, 30 seconds at 58 °C, 30 seconds at 72 °C), and 10 minutes at 72 °C. The LightCycler 480 II software was used to measure fluorescence and calculation of Ct-values (Roche, Software release 1.5.1.62 SP3). The fold change in expression of 16 genes were calculated by the 2^-ΔΔCt^ method using the geometric average of two reference genes (*PGK1* and *ACTB*) and comparing cells challenged with phages or *E. coli* ST-2064 relative to unchallenged cells (Supplementary Table S3).

### Animal housing, treatments and tissue collection

The animal experimental procedures were approved by the Danish Animal Experiments Inspectorate (license no. 2020-15-0201-00520). Five domestic sow-reared crossbred piglets (three male, two female) (Large white x Danish landrace x Duroc) with birthweight < 1000 grams were purchased from a Danish farm at two days-of-age. All animals received a broad-spectrum antibiotics cocktail consisting of amoxicillin-clavulanate (50 mg/kg/day), gentamicin (5 mg/kg/day), and metronidazole (20 mg/kg/day) to eliminate potential bacterial hosts for the phage. The antibiotics were administered through an oral feeding tube twice per day for two consecutive days. Approximately one hour after the first and third treatments, the phage Eco30 was administered through the oral feeding tube in a titer of 1 x 10^10^ PFU/kg bodyweight. The piglets were fed 20 ml/kg whole milk suspended from powder (Arla Pro, per 100 gram: 2110 kJ, 28 gram fat, 39 gram carbohydrate, 24 gram protein) every 3 hours between 6 AM and 9 PM, and 30 ml/kg at 12 AM. The pigs were weighed daily, and the clinical status, fecal output and consistency were assessed every 3 hours. On day three, the piglets were deeply anesthetized and euthanized by an intracardiac injection with pentobarbiturate (200 mg, Alfa Med). Thereafter, the contents of the small intestine and colon were collected in cryotubes. A section of the ileum was flushed with sterile saline, opened longitudinally, and mucosa was scraped off and collected in a cryotube. All cryotubes were snap-frozen in liquid nitrogen and stored at -80 °C until analysis.

### Sample analysis

The colon content was diluted 10-fold in 0.9% NaCl and cultured aerobically on MacConkey agar at 37 °C to estimate the density of aerobic Gram-negative bacteria. Another sample of the colon content and the small intestine content were directly transferred to Eppendorf tubes, whereas the small intestinal mucosa was first dissociated in 700 µl of SM buffer (GentleMACS Dissociator, Miltenyi Biotec) and then transferred to an Eppendorf tube. The tubes were centrifuged at 7.000 rpm for 5 min, the supernatant was centrifuged again, and passed through a 0.45 µm filter (MiniSart Syringe Filter, Sartorius). The phage titer was estimated in the filtrates of small intestine content and mucosa using the double agar overlay method as described . DNA was extracted from the filtrates of colon content, small intestine content and small intestine mucosa using QIAamp MinElute Virus Spin Kit (QIAGEN) according to the manufacturer’s instructions, followed by qPCR in duplicate reactions of 10 µl with primers specific for Eco30 (Supplementary Table S3), using the amplification protocol as described above. The difference in cycle threshold (Ct) between each sample and the averages were calculated and transformed using the 2^-ΔCt^ formula, and the relative phage quantity was calculated by dividing the individual values with the average.

### Statistical analysis

All statistical analyses were performed in RStudio (version 4.1.2), and data was visualized in GraphPad Prism (version 10.2.3). The fold change in gene expression was log2-transformed and compared using a linear model with Group and Replicate as fixed factors, followed by Tukey post-hoc test for comparison of groups. Unsupervised hierarchical clustering was performed with Ward’s minimum variance using R-packages “*factoextra*” and “*cluster*” and the difference in fold change between the two clusters was compared using an unpaired t-test. The small intestine phage titer was compared using a Wilcoxon test between locations (small intestine mucosa vs content), not assuming equal variance. Quantified phage DNA levels were compared pairwise between sample sites (small intestine mucosa vs small intestine content vs colon content) by a Wilcoxon test not assuming equal variance. Significance levels are: * p<0.05, ** p<0.01, *** p<0.001.

## Results

### Growth inhibitory effect of filtrates against bacteria from NEC-like mucosa

Mucosa collected from ileal biopsies with pronounced NEC-like pathology (extraluminal gas accumulations and necrosis) were collected from preterm piglets (n=11) and cultured aerobically on MacConkey agar to select for Gram-negative bacteria and obtain isolates belonging to *Enterobacteriaceae*. Nine mucosa samples were positive for bacterial growth, with a density ranging between 1.88 x 10^4^ CFU/ml to 2.23 x 10^8^ CFU/ml suspension (median 1.28 x 10^7^ CFU/ml). Based on the PFGE profiles, 16 unique bacterial isolates belonging to five species within the family *Enterobacteriaceae* were selected (Supplementary Table S1, Supplementary Figure S1).

Twenty-three donor fecal filtrates from healthy suckling piglets or sows (Supplementary Table S2; [14], [24]) were incubated individually with each bacterial isolate (Figure 1A). After 2 to 4 hours of incubation, six filtrates demonstrated growth inhibition in at least one replicate against three bacterial isolates , indicative of lytic phages in the filtrate specific for the investigated bacterial isolate (Figure 1B-D). The inhibition was either complete or partial, and the growth resumed with prolonged incubation, suggesting that the bacteria were able to develop resistance towards the phages. The bacterial isolates inhibited by the donor filtrates were analyzed by whole genome sequencing, and aligned with representative genomes to confirm the bacterial species identities as *Escherichia coli* ST-2064, *Escherichia coli* ST-10, and *Enterobacter hormaechi* ST-104 (Table 1). The sequenced isolates were further characterized by determination serotype and the presence of virulence and antibiotic resistance genes (Table 1).

**Figure 1.**
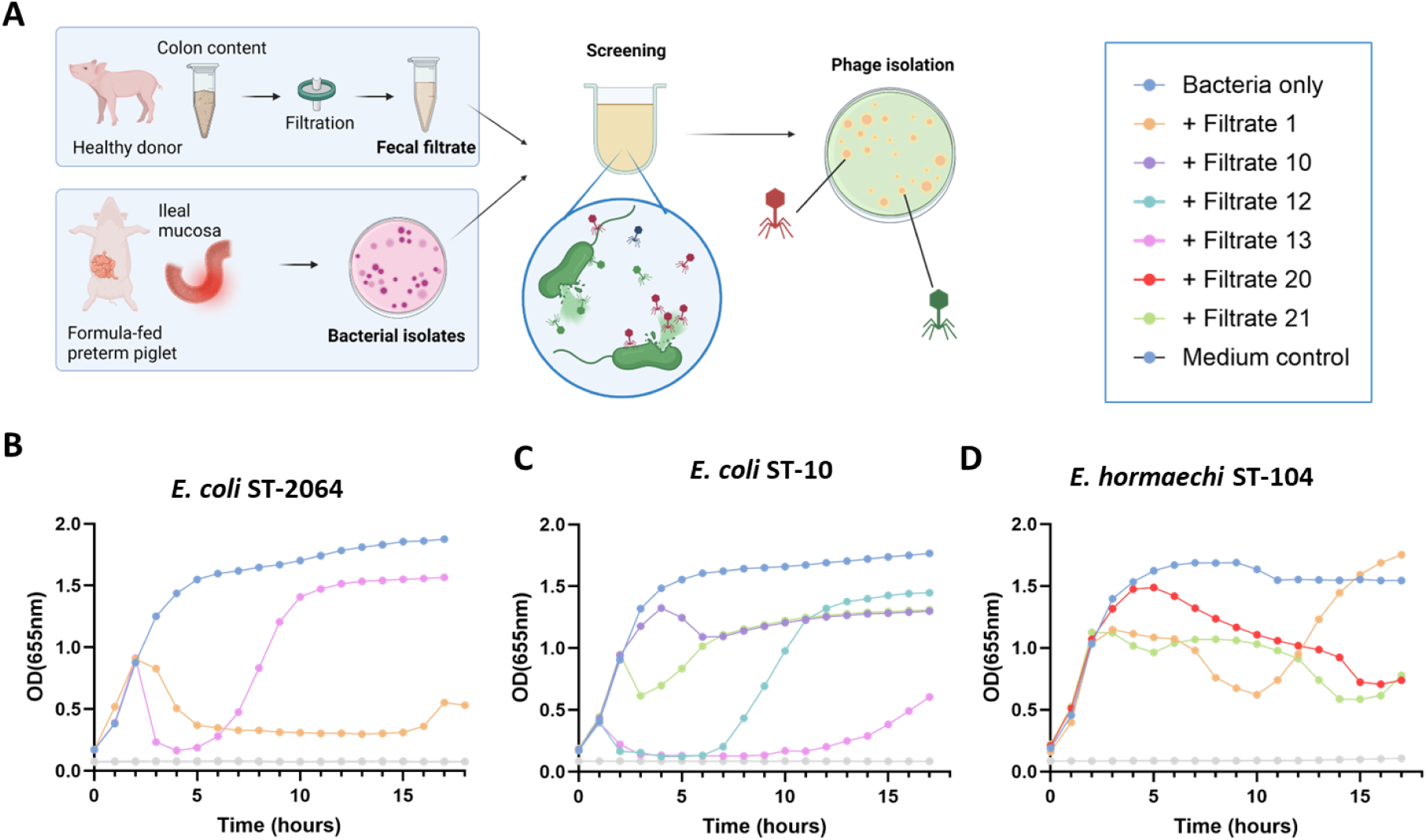
Six donor fecal filtrates inhibited growth of three bacterial strains isolated from NEC-like inflamed ileal mucosa. A) Overview of donor fecal filtrate preparation, culture of bacterial isolates from inflamed ileal mucosa, and screening for bacterial growth and isolation of phages. B)-D) Growth inhibition of *E. coli* ST-2064, *E. coli* ST-10, and *E. hormaechi* ST-104, respectively. Wells with inhibited bacterial growth is shown and represent one replicate per line.

**Table 1.**
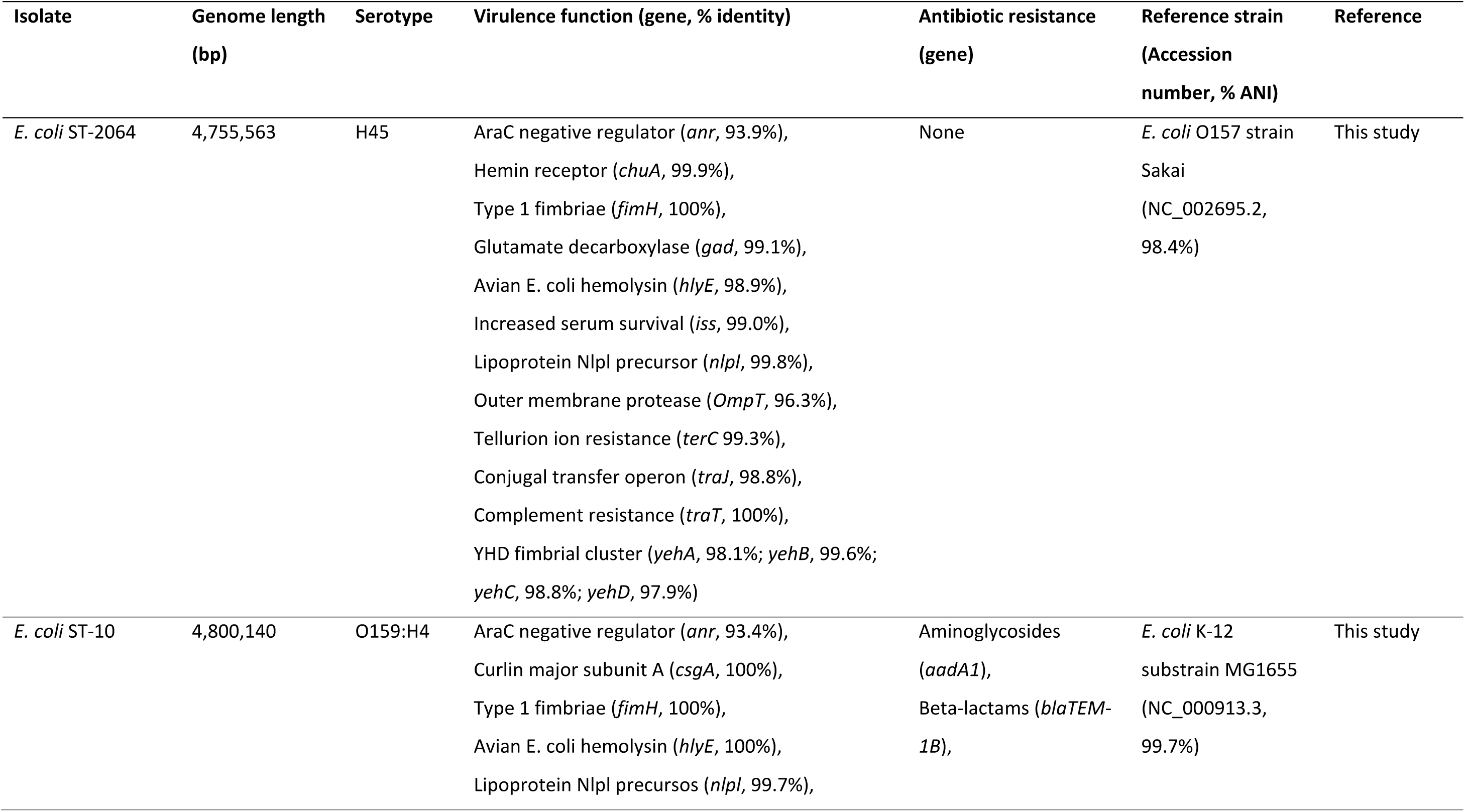
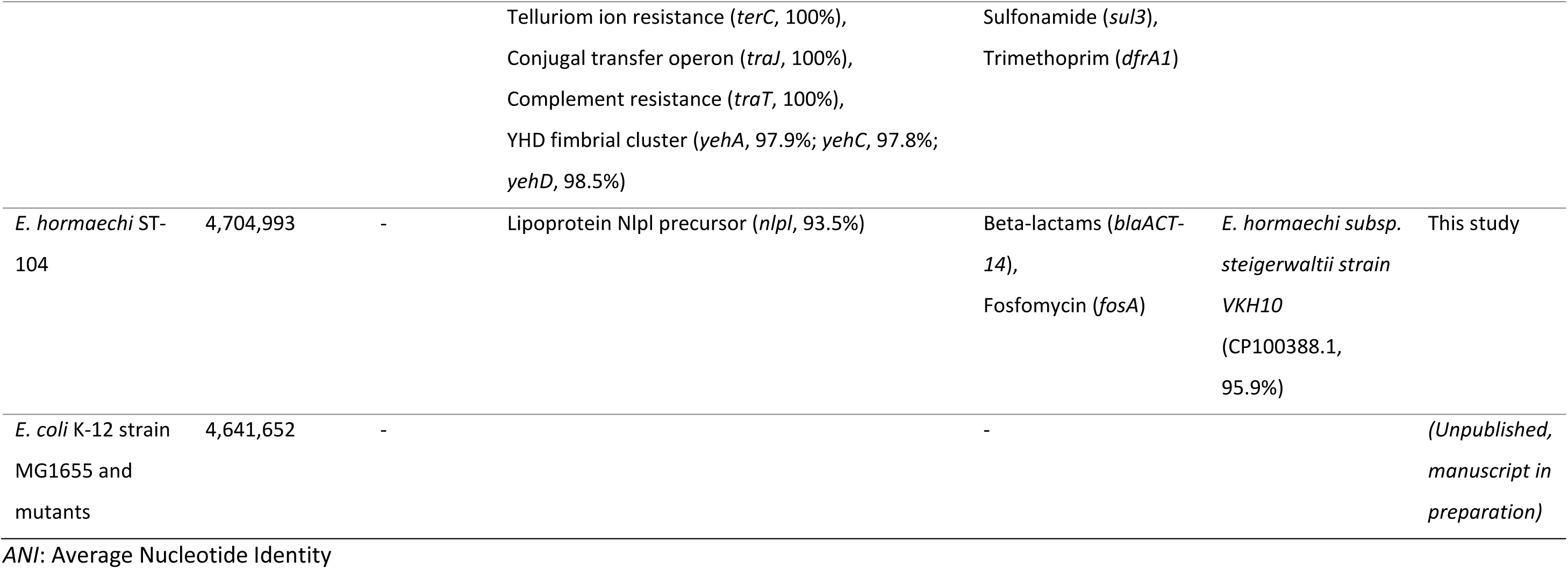
Bacterial strains used as bacteriophage hosts or to investigate the host binding structure.

| Isolate | Genome length<br>(bp) | Serotype | Virulence function (gene, % identity) | Antibiotic resistance<br>(gene) | Reference strain<br>(Accession<br>number, % ANI) | Reference |
| --- | --- | --- | --- | --- | --- | --- |
| *E. coli* ST-2064 | 4,755,563 | H45 | AraC negative regulator (*anr*, 93.9%),<br>Hemin receptor (*chuA*, 99.9%),<br>Type 1 fimbriae (*fimH*, 100%),<br>Glutamate decarboxylase (*gad*, 99.1%),<br>Avian *E. coli* hemolysin (*hlyE*, 98.9%),<br>Increased serum survival (*iss*, 99.0%),<br>Lipoprotein Nlpl precursor (*nlpl*, 99.8%),<br>Outer membrane protease (*OmpT*, 96.3%),<br>Tellurion ion resistance (*terC* 99.3%),<br>Conjugal transfer operon (*traJ*, 98.8%),<br>Complement resistance (*traT*, 100%),<br>YHD fimbrial cluster (*yehA*, 98.1%; *yehB*, 99.6%;<br>*yehC*, 98.8%; *yehD*, 97.9%) | None | *E. coli* O157 strain<br>Sakai<br>(NC_002695.2,<br>98.4%) | This study |
| *E. coli* ST-10 | 4,800,140 | O159:H4 | AraC negative regulator (*anr*, 93.4%),<br>Curlin major subunit A (*csgA*, 100%),<br>Type 1 fimbriae (*fimH*, 100%),<br>Avian *E. coli* hemolysin (*hlyE*, 100%),<br>Lipoprotein Nlpl precursors (*nlpl*, 99.7%), | Aminoglycosides<br>(*aadA1*),<br>Beta-lactams (*blaTEM-1B*), | *E. coli* K-12<br>substrain MG1655<br>(NC_000913.3,<br>99.7%) | This study |
|  |  |  | Telluriom ion resistance (*terC*, 100%),<br>Conjugal transfer operon (*traJ*, 100%),<br>Complement resistance (*traT*, 100%),<br>YHD fimbrial cluster (*yehA*, 97.9%; *yehC*, 97.8%;<br>*yehD*, 98.5%) | Sulfonamide (*sul3*),<br>Trimethoprim (*dfrA1*) |  |  |
| *E. hormaechi* ST-104 | 4,704,993 | - | Lipoprotein Nlpl precursor (*nlpl*, 93.5%) | Beta-lactams (*blaACT-14*),<br>Fosfomycin (*fosA*) | *E. hormaechi* subsp. *steigerwaltii* strain VKH10 (CP100388.1, 95.9%) | This study |
| *E. coli* K-12 strain MG1655 and mutants | 4,641,652 | - |  | - |  | (Unpublished, manuscript in preparation) |

ANI: Average Nucleotide Identity

### Establishing a phage consortium comprising members from three families

After the initial screening, we confirmed the presence of phages in the filtrates by plaque assay and isolated individual phages (Supplementary Figure S2). In total, we isolated 42 phages that infected either *E. coli* ST-2064 (n=9 phages), *E. coli* ST-10 (n=20 phages), or *E. hormaechi* ST-104 (n=13 phages). We excluded duplicate phages exhibiting the same plaque morphology, filtrate origin and bacterial host, and analyzed the remaining 21 phages by whole genome sequencing. The phage collection was further narrowed by genomic comparison and exclusion of identical phages. One phage contained a gene encoding an integrase and was also excluded.

The final collection consisted of ten strictly virulent phages with double-stranded DNA (dsDNA) genome, named according to published guidelines [34], and which will be referred to with their short names (Table 2). The phage genomes did not contain any genes associated with lysogeny, virulence or antibiotic resistance. Pairwise alignment of the phage genomes confirmed that while some shared high similarity, all phages were unique (Supplementary Figure S3). Notably, high sequence similarity (>90%) was observed between phages Eco27 and Eco30, between Eco09, Eco12 and Eco22, as well as between Ecl15 and Ecl18. Phage Ecl03 shared a similarity of 88.3% with Ecl15 and 87.1% with Ecl18, while phages Eco05 and Eco15 were the most distinct, sharing only 56% sequence similarity with each other and less than 15% with the other isolated phages.

**Table 2.**
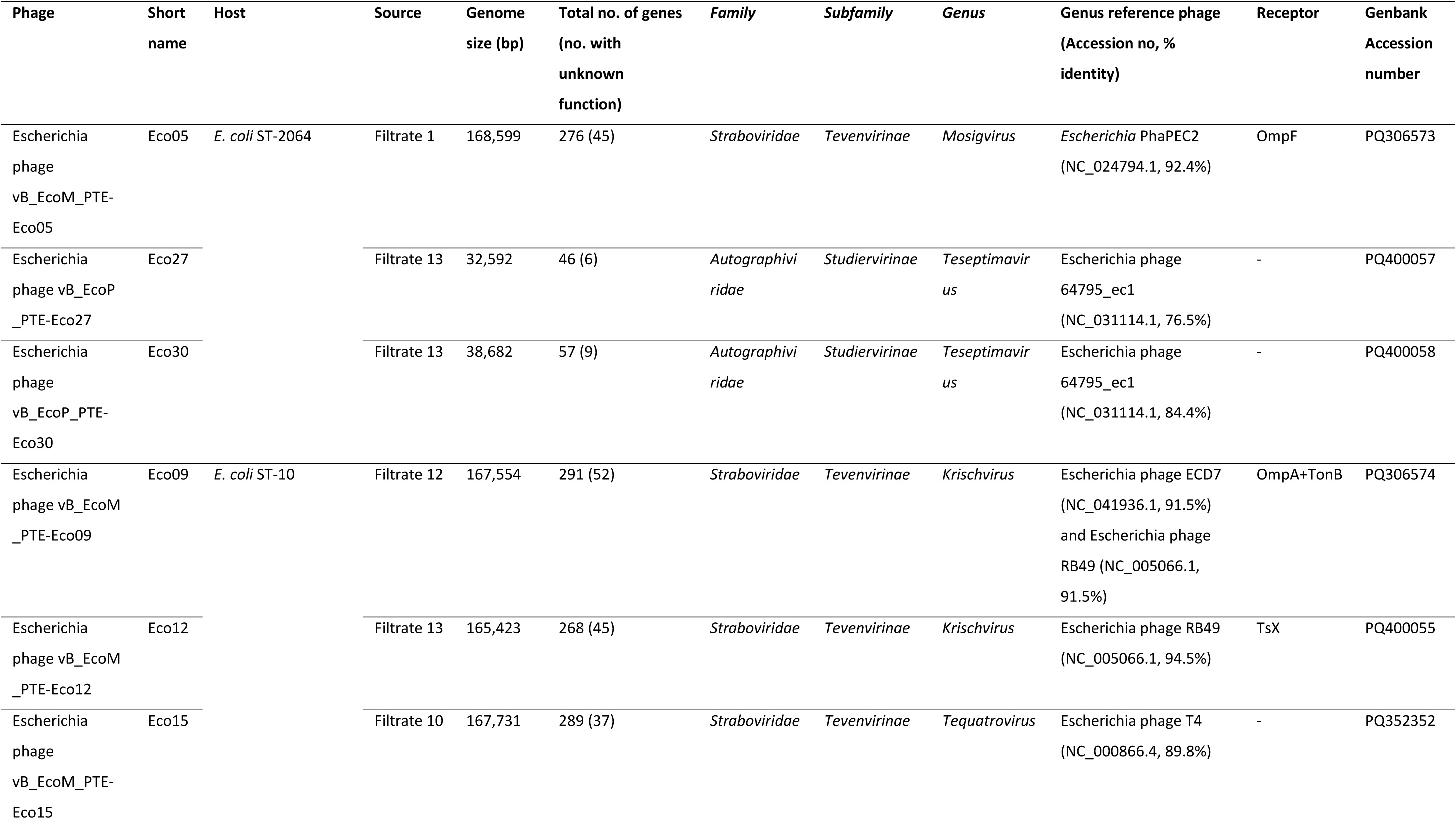
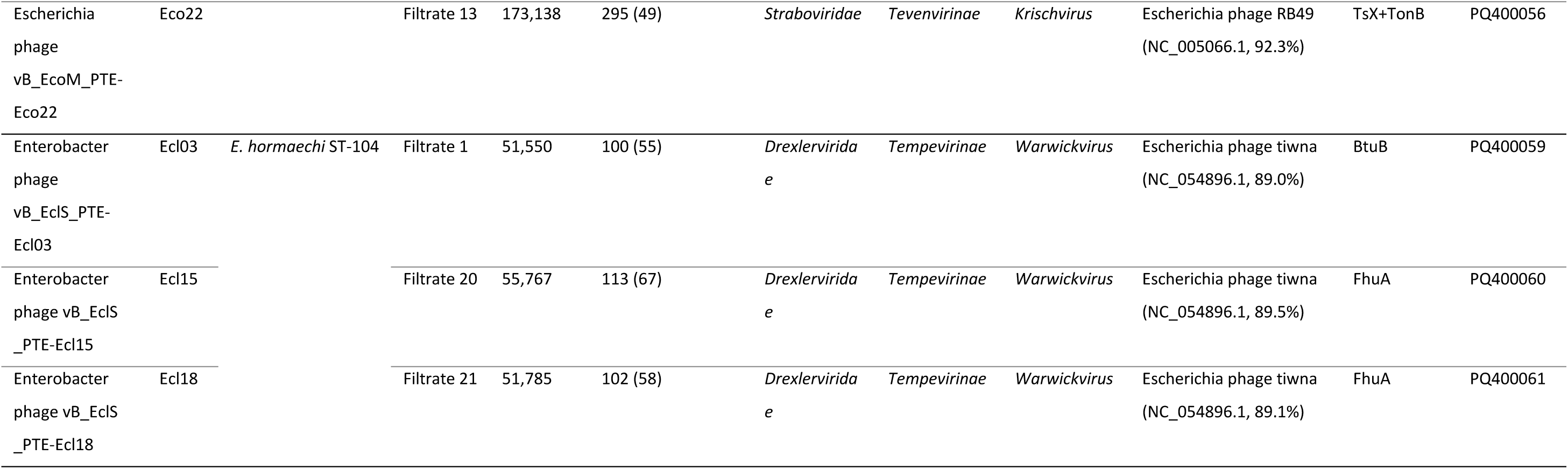
Bacteriophages isolated from donor fecal filtrates.

A BLAST search revealed high genomic similarity of the isolated phages (between 92% and 99%) to known phages in the *Caudoviricetes* order within the families *Straboviridae, Autographviridae* and *Drexlerviridae*. Phage taxonomy and phylogeny were assessed by genome comparison with reference phages for each genus within the subfamilies (Figure 2A, Table 2). Phage Eco05 showed the highest similarity to the *Escherichia* phage PhaPEC2, a reference for the *Mosigvirus* genus in the *Tevenvirinae* subfamily (*Straboviridae* family). The phages Eco09, Eco12 and Eco22 showed high genome similarity with reference phages for *Krischvirus*, also in the *Tevenvirinae* subfamily, while phage Eco15 showed high similarity to *Escherichia* phage T4, the reference phage for the *Tequatrovirus* genus in the *Tevenvirinae* subfamily. Phages Eco27 and Eco30 shared most similarity with the *Escherichia* phage 64795_ec1, reference phage of the *Teseptimavirus* genus in the *Studiervirinae* subfamily (*Autographviridae* family). Finally, phages Ecl03, Ecl15 and Ecl18 exhibited the highest genome similarity to *Escherichia* phage tiwna, reference phage for the *Warwickvirus* genus in the *Tempevirinae* subfamily (*Drexlerviridae* family).

**Figure 2.**
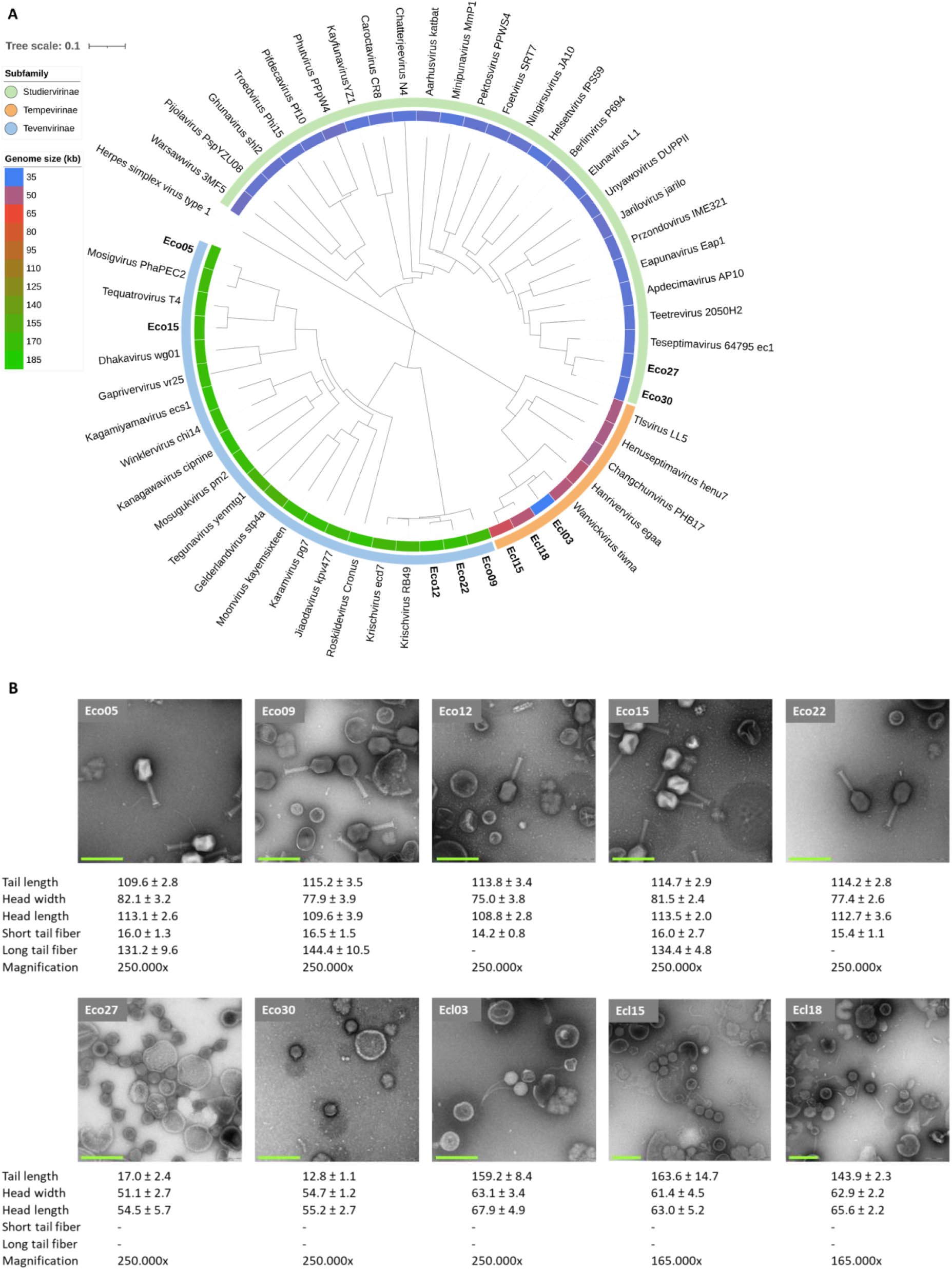
The isolated phages belong to three subfamilies and display different morphology. A) Phylogenetic tree of isolated and representative phages from each genus in the subfamilies prepared from the nucleotide distance algorithm, phages isolated in this paper are marked in bold. B) TEM images of negative stained phages. Dimensions of tail, head, short and long tail fibers were measured on five phage particles. All measurements are in nm, shown as mean and standard deviation, scale: 200 nm.

The morphology of each phage was characterized by transmission electron microscopy (TEM) (Figure 2B, 2C). All phages in the *Tevenvirinae* subfamily (Eco05, Eco09, Eco12, Eco15, and Eco22) displayed myovirus morphology, featuring an elongated icosahedral head and a long contractile tail, although their dimensions varied slightly. Phages Eco05, Eco09 and Eco15 additionally carried long tail fibers. In contrast, phages Eco27 and Eco30 displayed a podovirus morphology characterized by an isometric head and short tail. Lastly, phages Ecl03, Ecl15, and Ecl18 showed a siphovirus morphology featuring a rounded head and a long, flexible, non-contractile tail.

### Different host receptors utilized by the phages in the consortium

Receptor binding is an essential step of phage infection. To determine the structural basis for phage-host binding, we evaluated the infective capacity of the isolated phages against the laboratory strain *Escherichia coli* MG1655 and its derived mutants deficient in known phage receptors (Δ*btuB*, Δ*fadL*, Δ*fepA*, Δf*huA*, Δ*lamB*, Δ*ompA*, Δ*ompC*, Δ*ompF*, Δ*ompU*, Δ*tolC*, Δ*tonB*, Δ*tsX)*. The phages Eco15, Eco27, and Eco30 were unable to infect *E. coli* MG1655 WT and were not evaluated on the receptor mutants. The remaining phages exhibited reduced or inhibited infection of various mutant strains, indicating different structures necessary for phage infection. Specifically, phages Eco05 and Eco09 respectively showed decreased infectivity of the strains MG1655 Δ*ompA* and MG1655 Δ*ompC,* deficient in different outer membrane proteins (Figure 3A). Phages Eco12 and Eco22 demonstrated reduced infectivity in the mutant MG1655 Δ*tsX*, encoding a nucleoside transporter (Figure 3B).

**Figure 3.**
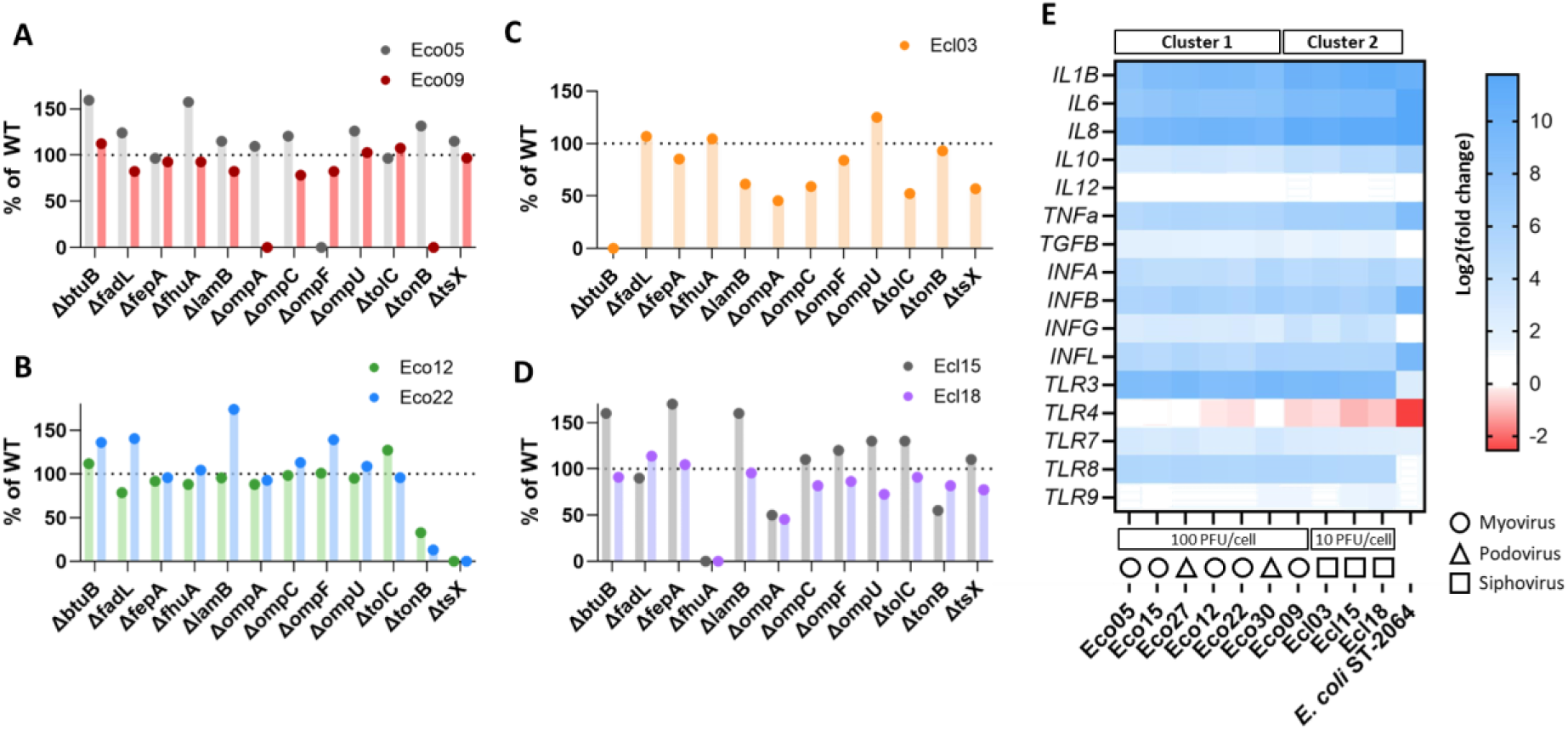
The phages utilize various surface structures for host binding and induce different levels of immune gene expression in monocyte-derived macrophages. A-D) Phage infectivity on *E. coli* MG1655 receptor mutants as a percentage of the infective titer on the WT strain. BtuB: Vitamin B12-transporter. FadL: Long-chain fatty acid transporter. FepA: Ferric enterobactin transporter. FhuA: Ferrichrome transporter. LamB: Malto-oligosaccharide transporter. OmpA: Outer membrane protein A. OmpC: Outer membrane protein C. OmpF: Outer membrane protein F. OmpU: Outer membrane protein U. TolC: Outer membrane transporter. TonB: Energy transducer. TsX: Nucleoside transporter. E) Cell gene expression shown as log2(fold change in gene expression) relative to unchallenged cells. n=3 biological replicates. All involved genes, with exception for IL-12 in cells challenged with Eco05, and TLR4 for all phages, showed increased expression relative to unchallenged cells.

Furthermore, phage Ecl03 was completely unable to infect the MG1655 Δ*btuB* mutant strain (Figure 3C), while phages Ecl15 and Ecl18 were unable to infect MG1655 Δ*fhuA* (Figure 3D). The phages Eco09 and Eco22 additionally exhibited reduced infectivity of the MG1655 Δ*tonB* strain, lacking a key structure that is coupled with the activity of outer membrane proteins [35]. Collectively, the phages in our selected consortium utilized a variety of host receptors, allowing them to potentially infect a broader range of hosts that present these structures on their surface.

### Consistent immunogenicity of phages in the consortium

Phages can induce a strong immune response in mammalian cells [36]. We explored the immune response induced by the isolated phages in the THP-1 cell line of monocyte-derived macrophages, compared with sterile buffer or *E. coli* ST-2064 as reference for Gram-negative bacterial stimulation. All ten phages increased expression of both pro-and anti-inflammatory genes, as well as the expression of nucleotide-specific receptors TLR3, TLR7, TLR8 and TLR9, but to varying extents (Supplementary Table S4). Based on the cellular responses, we classified the phages into two distinct clusters through unsupervised clustering analysis, and observed distinct changes in the cellular gene expression for each cluster and compared to *E. coli* (Figure 3E). Phages with myovirus morphology were represented in both clusters, while podoviruses were exclusively present in cluster 1, and siphoviruses were restricted to cluster 2. In both clusters, the expression levels of genes *IL1B, IL6, IL8, IL10, IL12, TNFa, TGFB, INFG, TLR7,* and *TLR9* were elevated compared with unchallenged cells, although to a significantly higher level in cluster 2. Expression levels of the genes *INFB, INFL, TLR3,* and *TLR8* were similar between the two clusters, although all were still elevated compared to unchallenged cells. Notably, the expression of *TLR4* was downregulated in response to *E. coli* and selected phages (Eco22, Eco09, Ecl03, Ecl15, and Ecl18). In contrast, all phages induced a significant increase in the expression of *TLR3* and *TLR8*, along with a modest increase in *TLR7* and *TLR9*. Cells incubated with *E. coli* ST-2064 showed a marked change in gene expression levels compared to those co-cultured with phages. Specifically, the increases in *IL6, IL8, IL10, TNFa, INFB,* and *INFL* expression levels were greater for cells co-cultured with *E. coli* ST-2064 than in cells co-cultured with phages, while the *TGFB, INFG, TLR3, TLR4,* and *TLR8* expression levels were lower.

These observations support the findings that phages can evoke an immune response in phagocytic cells, likely independent on the phage morphology and to a different extent than Gram-negative bacteria.

### High-dose phage therapy in newborn piglets and detection of infective phages in the gut

As the final investigation, we tested the clinical responses and fate of a selected phage in newborn piglets. We selected Eco30 from the phage consortium, as it was readily propagated to a high titer and formed large, clearly distinct plaques. The phage was administered at a high dose of 10^10^ PFU per kg body weight to five two-day-old piglets receiving broad-spectrum antibiotics to deplete potential phage host bacteria (Figure 4A). All animals had natural body weight increments and transient, mild diarrheal episodes, as often seen after a change of environment, and no single animal showed adverse events attributed to phage treatment. At euthanasia, the Gram-negative bacterial density in colon content ranged between 3.8 x 10^3^ to 9.5 x 10^5^ CFU/ml colon content (median 6.9 x 10^3^ CFU/ml). The infective phage titer was assessed in samples of ileal luminal content and washed ileal mucosa to assess the propensity for mucosa association. Plaques with the expected morphology for phage Eco30 (Supplementary Figure S4B) were observed at double agar overlay with the host strain and confirmed that the administered phage remained infective in both the mucosa and content at comparable levels (p=0.6) (Figure 4B). The quantification of phage-specific DNA signatures further supported this finding, indicating that the phage Eco30 was present in similar quantities across the two sites (p=0.37) (Figure 4C). Nevertheless, the relative phage DNA quantity in the colon content was slightly higher than in the ileal mucosa and only trending higher than in the ileal content.

**Figure 4.**
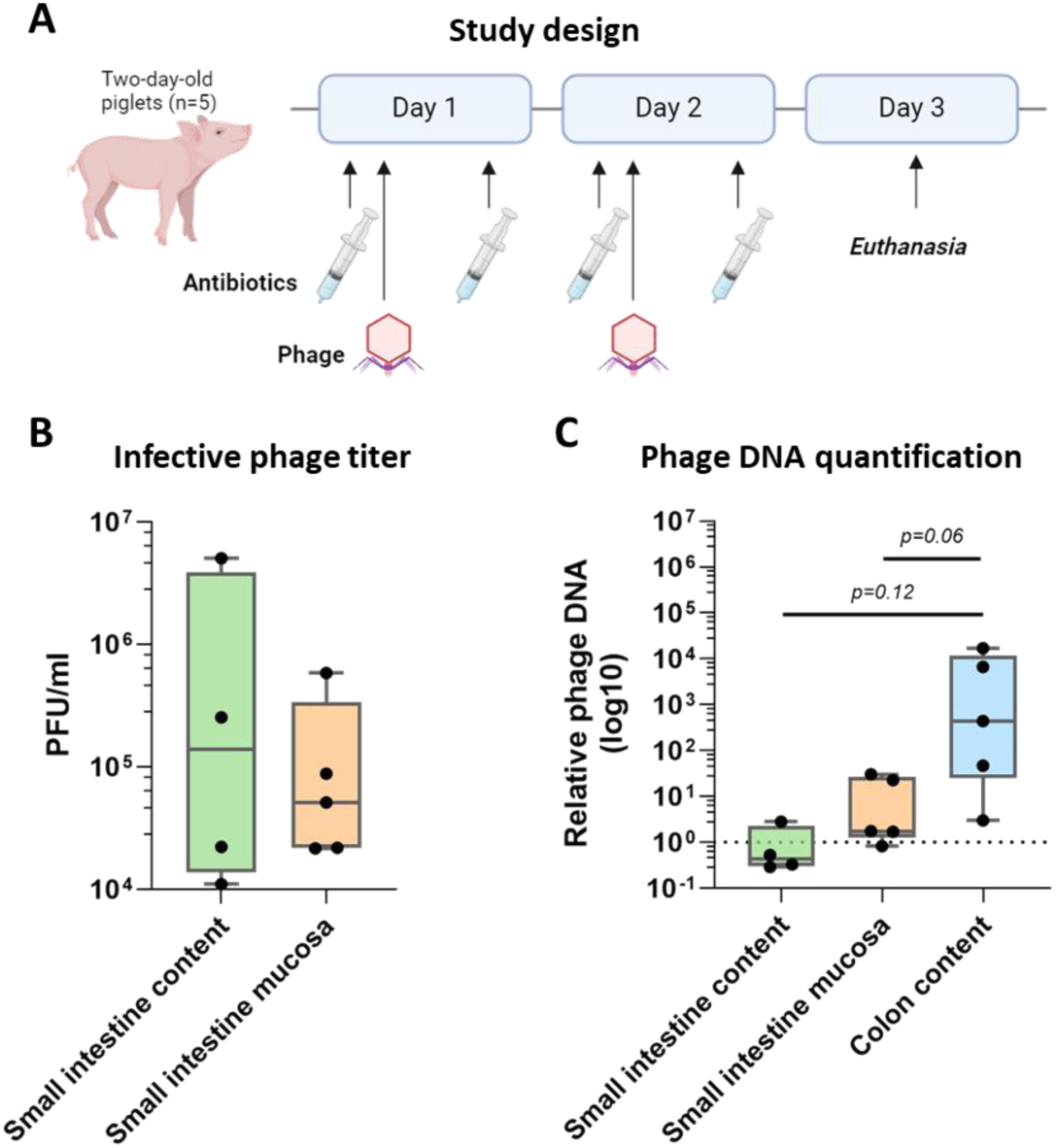
The fate of orally administered phage in newborn piglets. A) Study overview with arrows indicating the administration of antibiotics or phages. B) Infective phage density in the small intestine content and mucosa. C) Quantification of phage DNA in small intestine content, small intestine mucosa, and colon content. n=4-5 animals

In summary, we detected the phage as infective particles or quantifiable DNA signatures following oral administration in newborn piglets, suggesting that it survives and remains active in gastrointestinal conditions. The phage administration appeared to be safe in healthy newborn piglets, with no obvious adverse effects observed.

## Discussion

In this study, we isolated and characterized a collection of ten unique lytic phages that infect *Enterobacteriaceae* associated within the gut mucosa of newborn piglets exhibiting NEC-like pathology. The phage collection exclusively includes virulent phages to mitigate the risk of gene transfer between hosts during infection associated with temperate phages [37]. Additionally, the isolated phages utilized various structures to bind to their bacterial hosts and initiate infection, consistent with the structures identified as important for other similar phages [38]. The three siphovirus phages (Ecl03, Ecl15, Ecl18) were completely unable to infect mutant strains of *E. coli* MG1655 that lacked the ferric siderophore FhuA, or the vitamin B12 transporter BtuB, both known phage targets [39]. A prior large-scale analysis of over 60 phages identified these structures as essential for infection of siphovirus belonging to the *Warwickvirus* genus, in accordance with the classification of the phages in this study [38]. The phage belonging to the *Mosigvirus* genus (Eco05) relied on OmpF for infection, while the phages in the *Krischvirus* genus (Eco09, Eco12 and Eco22) depended on OmpA or the nucleoside transporter TsX. All three outer membrane proteins are characteristic receptors for phages within these genera [38]. Two phages demonstrated reduced infectivity in the strain lacking TonB, an essential mediator of substrate transport via outer membrane proteins [35]. It is possible that the phages might not utilize TonB directly, but rather that its absence affects their primary targets. Among the isolated phages, three (Eco15, Eco27 and Eco30) were unable to infect the *E. coli* MG1655 WT strain, and their binding structures could not be determined using the receptor-deficient mutant strains. Given that *E. coli* MG1655 lacks the O-antigen component on the lipopolysaccharide (LPS) layer, these phages may rely on this component for binding [40] or may be unable to infect due to other bacterial resistance mechanisms.

In the *in vitro* co-culture, we found that all ten phages elicited immune responses to varying extents, and that the cellular response was independent of the phage titer. As all the siphovirus phages along with one myovirus phage clustered with increased gene expression, we speculate that the intrinsic structures and properties of phages are critical determinants of immunogenicity. It is well established that phages can be recognized by the mammalian immune system and influence inflammatory responses, cellular growth and bacterial clearance, but the specific structures responsible for the differential immune responses remain unclear [36], [41], [42], [43], [44], [45].

These findings suggest that the immunological response to phages is contingent on disease and experimental parameters, thereby necessitating careful evaluation before considering them for therapeutic applications. A previous study indicated a reduction in *TLR4* expression in human blood-derived monocytes co-cultured with a podovirus phage [36]. In contrast, our study revealed that five of the ten phages significantly decreased the *TLR4* expression relative to unchallenged cells.

Conversely, all phages increased the expression of *TLR3*, *TLR7* and *TLR8*, which respond to viral RNA. This outcome is consistent with another study, which reported increased TLR3 signaling and inhibited TNF-α production in murine phagocytes co-cultured with a dsDNA phage [44]. Given that the isolated phages contained dsDNA genomes, it is plausible that the elevated *TLR3* expression could be a result from downstream signaling in phage-stimulated cells. Alternatively, the increase in expression might be partially attributed to the presence of phage RNA in the administered phage preparations, or the uptake of phages into immune cells leading to phage RNA production [44].

Recent advancements in microbiome-targeting treatments involve fecal microbiota transplantation (FMT) from healthy donors to alleviate gut diseases associated with inflammation [46]. While FMT exhibited reduced intestinal inflammation and the incidence of NEC in preterm piglets, it also carried a high risk of sepsis-related mortality [47]. Sterile FFT may offer a safer alternative, as exemplified by promising outcomes in preterm piglets and in adult patients with recurrent *Clostridioides difficile* infection [14], [48]. The effects of FFT are likely attributed to the activity of phages in the fecal virome [15]. While the impact of FMT and FFT on alleviating gut bacterial dysbiosis is evident, the variation in microbial composition of donor materials could limit their clinical applications [46], [49].

Administration of a pre-defined phage consortium that specifically targets bacteria associated with gut inflammation would provide a reliable treatment. In the current study, we demonstrated retained presence and infectivity of the administered phage in the gut content and mucosa of newborn pigs. The lower phage titer in the small intestinal content could indicate that the phages are merely passing through the small intestine and instead adhere to the mucosal lining or accumulate in the colon. These and other phages may be combined in future studies to test their efficacy against NEC and other intestinal diseases in newborns.

We acknowledge the limitations of this study. Although we screened a large panel of bacteria and fecal filtrates, only limited filtrates exhibited growth inhibition. Further, it would be advantageous to culture bacteria associated with the colonic mucosa in addition to ileal mucosa, as colon pathology is more frequently observed in the piglet model of NEC [50]. While human infants developing NEC often harbor increased abundances of *Pseudomonadota* including *Enterobacteriaceae* [6], [7], [51], the preterm piglets developing NEC are also frequently linked with increased mucosal abundances of *Clostridiales* [52], [53]. However, the previous finding of reduced abundance of *Proteobacteria* in the ileal mucosa of piglets not developing NEC [14] prompted us to investigate these.

In conclusion, we have demonstrated the feasibility of isolating phages targeting bacteria associated with NEC-like pathology and inflammation. Further, we have proven their immunogenicity *in vitro*, as well as their presence and infectivity in the gut following oral administration into newborn piglets.

Future studies could explore the dissemination of other phages, and *in vivo* administration of phage consortia may further contribute to the advancement of phage therapy for newborn gut diseases such as NEC.

## Acknowledgements

Data for phage genome sequencing was generated through accessing research infrastructure at University of Copenhagen, including FOODHAY (Food and Health Open Innovation Laboratory, Danish Roadmap for Research Infrastructure). TEM analysis was performed with microscopes and assistance at the Core Facility for Integrated Microscopy, Faculty of Health and Medical Sciences, University of Copenhagen. The *E. coli* MG1655 WT and receptor mutants were supplied from Veronika Lutz, Section for Food Safety and Zoonoses, University of Copenhagen. The THP-1 monocytic cell line was obtained from Professor John Elmerdahl Olsen, Section for Veterinary Clinical Microbiology, University of Copenhagen.

