## Supplemental Figures and Tables for "Characterization of a virulent bacteriophage consortium targeting Enterobacteriaceae from inflamed preterm gut mucosa"

### Supplementary Figures

#### Supplementary Figure S1

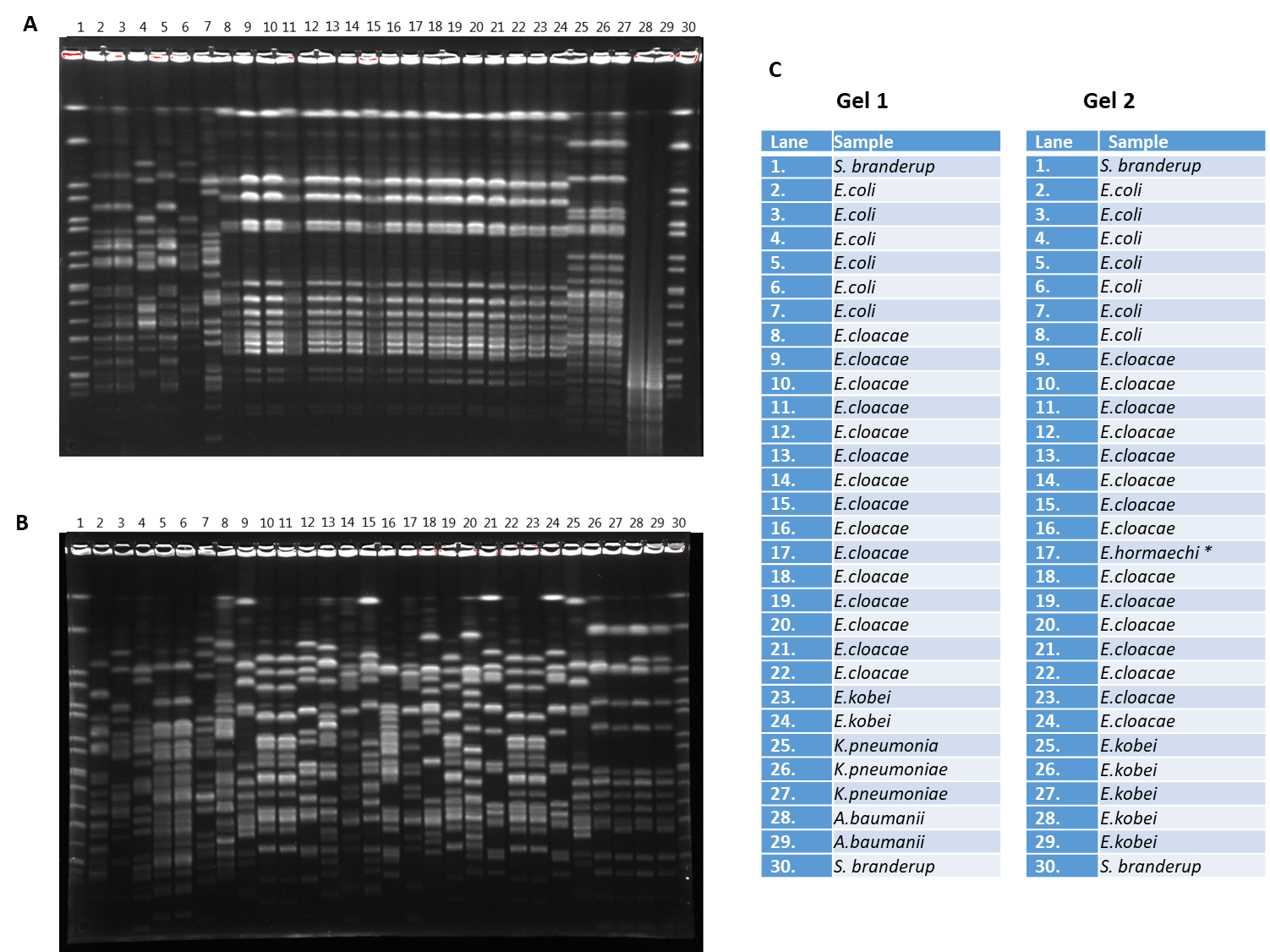

**Supplementary Figure S1.** Pulse-Field Gel Electrophoresis (PFGE) profiling of bacteria isolated from inflamed ileal mucosa to select unique isolates.
A) Gel 1. B) Gel 2. C) Overview of samples.
The *Salmonella* serovar *Branderup* digested with restriction enzyme XbaI was used as a size marker (located on lanes 1 and 30 on both Gel 1 and Gel 2).
* The isolate was initially identified as *E. cloacae* by MALDI-TOF MS, but whole genome sequencing confirmed its identity as *E. hormaechi.*

#### Supplementary Figure S2

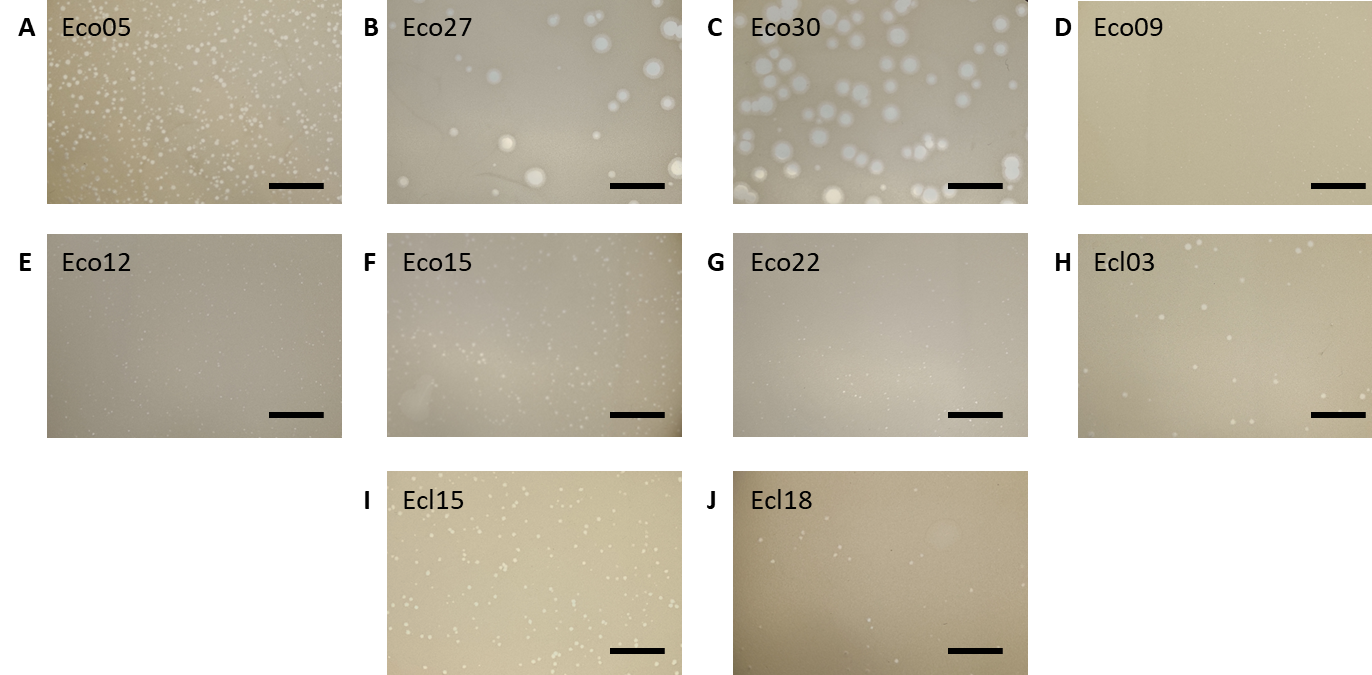

**Supplementary Figure S2.** Plaques of the ten selected phages in the consortium isolated from donor fecal filtrates, infecting three bacterial strains from inflamed ileal mucosa. Scale: 1 cm

#### Supplementary Figure S3

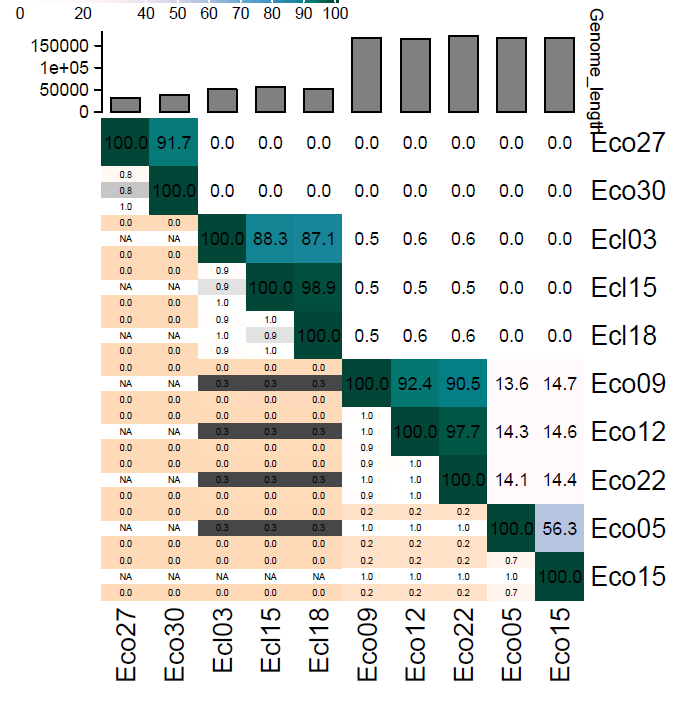

**Supplementary Figure S3**. Comparison of the genome similarity of the ten selected phages in the consortium isolated from donor fecal filtrates, and infecting three bacterial strains from inflamed ileal mucosa. Alignment of the genomes performed with VIRIDIC. Color indicates genome similarity in percentage.

#### Supplementary Figure S4

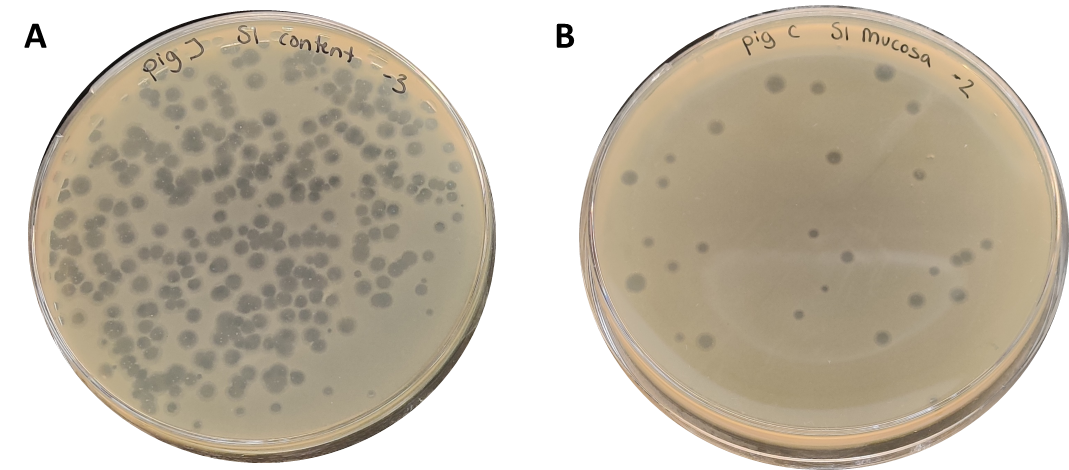

**Supplementary Figure S4.** Phages remain infective during upper gastrointestinal transit in preterm piglets administered a phage cocktail of a selected phage isolated from donor fecal filtrates, and infecting an *E. coli* isolated from inflamed ileal mucosa
A) Plaques from small intestine content, 1000x dilution. B) Plaques from small intestine mucosa, 100x dilution. Diameter of plate: 9 cm.

### Supplementary Tables

#### Supplementary Table S1: Bacterial isolates from inflamed ileal mucosa of preterm piglets sampled on day five of life

| **Origin** | **Species** | **Colony appearance**  **(color and diameter)** | **Growth inhibited by filtrates** |
| --- | --- | --- | --- |
| Piglet 1 | *Enterobacter cloacae* | Pink, 3-4 mm | No |
| Piglet 2 | *Enterobacter cloacae* | Pink, 3-4 mm | No |
| Piglet 7 | *Enterobacter cloacae* | Pink, 5 mm | No |
| Piglet 15 | *Enterobacter cloacae* | Transparent, 4-5 mm | No |
| Piglet 15 | *Enterobacter cloacae* | Transparent, 4-5 mm | No |
| Piglet 15 | *Enterobacter cloacae* | Pink, 4-5 mm | No |
| Piglet 16 | *Enterobacter cloacae* | Transparent, 4-5 mm | No |
| Piglet 16 | *Enterobacter cloacae* | Transparent, 4-5 mm | No |
| Piglet 17 | *Enterobacter cloacae* | Transparent, 5 mm | No |
| Piglet 17 | *Enterobacter cloacae* | Pink, 4-5 mm | No |
| Piglet 17 | *Enterobacter cloacae* | Transparent, 2 mm | No |
| Piglet 16 | *Enterobacter hormaechi* | Pink, 4-5 mm | Yes |
| Piglet 16 | *Enterobacter kobei* | Pink, 2-3 mm | No |
| Piglet 5 | *Escherichia coli* | Pink/purple, 5 mm | Yes |
| Piglet 5 | *Escherichia coli* | Pink/purple, 2 mm | Yes |
| Piglet 7 | *Klebsiella pneumoniae* | Pink, 5 mm | No |

#### Supplementary Table S2: Donor fecal filtrates and origin

| **Filtrate** | **Donor** | **Reference** |
| --- | --- | --- |
| Filtrate 1 | Pooled colon content from four piglets | (Brunse *et al.*, 2022) |
| Filtrate 2 | Colon content from 10-day old piglet | - |
| Filtrate 3 | Colon content from 10-day old piglet | - |
| Filtrate 4 | Colon content from 10-day old piglet | - |
| Filtrate 5 | Colon content from 10-day old piglet | - |
| Filtrate 6 | Colon content from 8-day old piglet | - |
| Filtrate 7 | Colon content from 11-day old piglet | - |
| Filtrate 8 | Colon content from 12-day old piglet | - |
| Filtrate 9 | Colon content from 10-day old piglet | - |
| Filtrate 10 | Colon content from 9-day old piglet | - |
| Filtrate 11 | Colon content from 11-day old piglet | - |
| Filtrate 12 | Colon content from 11-day old piglet | - |
| Filtrate 13 | Colon content from 11-day old piglet | - |
| Filtrate 14 | Colon content from 10-day old piglet | - |
| Filtrate 15 | Colon content from 12-day old piglet | - |
| Filtrate 16 | Colon content from 11-day old piglet | - |
| Filtrate 17 | Colon content from 10-day old piglet | - |
| Filtrate 18 | Colon content from 11-day old piglet | - |
| Filtrate 19 | Pool of feces from piglets | (Gambino *et al.*, 2024) |
| Filtrate 20 | Pool of feces from sows with suckling piglets | (Gambino *et al.*, 2024) |
| Filtrate 21 | Pool of feces from sows with suckling piglets | (Gambino *et al.*, 2024) |
| Filtrate 22 | Pool of feces from weaned piglets | (Gambino *et al.*, 2024) |
| Filtrate 23 | Pool of feces from 30 kg piglets treated with antibiotics | (Gambino *et al.*, 2024) |

All piglets included as donors were healthy and born at term

#### Supplementary Table S3: Primer sequences

| Gene | Forward primer | Reverse primer | Reference |
| --- | --- | --- | --- |
| *IL1B* | GGCCACATTTGGTTCTAAGAAA | TAAATAGGGAAGCGGTTGCTC | (Van Belleghem *et al.*, 2017) |
| *IL6* | GGTACATCCTCGACGGCATC | GCCTCTTTGCTGCTTTCACAC | (Van Belleghem *et al.*, 2017) |
| *IL8* | AGTTTTTGAAGAGGGCTGAGAAT | GCTTGAAGTTTCACTGGCATCT | (Kawarizadeh *et al.*, 2021) |
| *IL10* | CAACCTGCCTAACATGCTTCGAGAT | CCTCCAGCAAGGACTCCTTTAACAA | (Li *et al.*, 2022) |
| *IL12* | GCGGAGCTGCTACACTCTCT | GGTGGGTCAGGTTTGATGAT | (Zhang *et al.*, 2016) |
| *TNFa* | CCCAGGGACCTCTCTCTAATC | ATGGGCTACAGGCTTGTCACT | (Van Belleghem *et al.*, 2017) |
| *TGFB* | GAAGGGAGACAATCGCTTTAGC | TGTAGACTCCTTCCCGGTTGAG | (Van Belleghem *et al.*, 2017) |
| *INFA* | ATTTCTGCTCTGACAACCTC | TGACAGAGACTCCCCTGATG | (Chen *et al.*, 2012) |
| *IFNB* | AAACTCATGAGCAGTCTGCA | AGGAGATCTTCAGTTTCGGAGG | (Richtsteiger *et al.*, 2003) |
| *IFNG* | GACCAGAGCATCCAAAAGAGT | ATTGCTTTGCGTTGGACATTC | (Meenakshi *et al.*, 2016) |
| *IFNL* | GGACGCCTTGGAAGAGTCACT | AGAAGCCTCAGGTCCCAATTC | (Slater *et al.*, 2010) |
| *TLR3* | AACTCCATCTCATGTCCAACTCAA | GATGACAAGCCATTATGAGACAGATC | (Månsson *et al.*, 2010) |
| *TLR4* | CAGAGTTGCTTTCAATGGCATC | AGACTGTAATCAAGAACCTGGAGG | (Tedesco *et al.*, 2018) |
| *TLR7* | GGAGGTATTCCCACGAACACC | TGACCCCAGTGGAATAGGTACAC | (Vreća *et al.*, 2018) |
| *TLR8* | AACTTTCTATGATGCTTACATTTCTTATGAC | GGTGGTAGCGCAGCTCATTT | (Li *et al.*, 2013) |
| *TLR9* | GTCCAGCACTCGGAGGTTTC | TGGTGTTGAAGGACAGTTCTCTCT | (Mortezagholi *et al.*, 2017) |
| *ACTB* | AAGGTGACAGCAGTCGGTT | TGTGTGGACTTGGGAGAGG | (Wei *et al.*, 2023) |
| *PGK1* | CAAGAAGTATGCTGAGGCTGTCA | CAAATACCCCCACAGGACCAT | (Falkenberg *et al.*, 2011) |
| Phage Eco30 | CAAGGCGTCTTTAGGATGGT | TCCCTATCTGTTCAGCCTCC | This study |

#### Supplementary Table S4: Fold change gene expression in cells challenged with individual phages isolated from donor fecal filtrates or *E. coli* ST-2064 isolated from inflamed ileal mucosa, relative to unchallenged cells.

|  | **Eco05** | **Eco09** | **Eco12** | **Eco15** | **Eco22** | **Eco27** | **Eco30** | **Ecl03** | **Ecl15** | **Ecl18** | ***E. coli***  **ST-2064** |
| --- | --- | --- | --- | --- | --- | --- | --- | --- | --- | --- | --- |
| *IL1B* | 254.0 ± 68.5 | 1320.3 ± 372.0 | 646.9 ± 206.5 | 456.4 ± 123.3 | 605.2 ± 168.7 | 493.7 ± 114.7 | 444.4 ± 14.7 | 1118.5 ±  284.4 | 1823.8 ±  851.0 | 1005.2 ±  828.6 | 1542.5 ± 686.7 |
| *IL6* | 152.1 ± 26.5 | 560.1 ± 127.0 | 243.4 ± 45.6 | 299.8 ± 63.0 | 242.1 ± 39.9 | 291.1 ± 67.0 | 294.4 ± 49.0 | 494.5 ± 44.2 | 593.5 ± 91.9 | 612.6 ± 93.7 | 3540.8 ± 1748.1 |
| *IL8* | 554.3 ± 137.2 | 2206.5 ± 704.7 | 1101.1 ± 258.0 | 786.4 ± 249.5 | 1016.2 ± 210.8 | 897.8 ± 236.8 | 806.0 ± 197.5 | 1791.4 ± 447.2 | 2574.3 ± 1035.8 | 2444.4 ± 695.9 | 3633.7 ± 1285.4 |
| *IL10* | 8.4 ± 1.4 | 18.6 ± 1.8 | 9.6 ± 0.4 | 8.4 ± 0.7 | 8.1 ± 0.8 | 11.2 ± 1.9 | 10.2 ± 1.3 | 17.5 ± 1.2 | 30.6 ± 4.1 | 29.5 ± 5.4 | 85.6 ± 29.8 |
| *IL12* | 1.4 ± 0.1 | 2.3 ± 0.2 | 1.7 ± 0.1 | 1.5 ± 0.2 | 1.6 ± 0.3 | 1.6 ± 0.1 | 1.6 ± 0.3 | 1.8 ± 0.1 | 1.9 ± 0.2 | 2.1 ± 0.5 | 1.8 ± 0.02 |
| *INFA* | 29.2 ± 13.9 | 27.6 ± 10.6 | 27.2 ± 18.4 | 23.4 ± 12.2 | 18.6 ± 3.7 | 23.7 ± 12.3 | 47.1 ± 30.8 | 36.8 ± 19.6 | 33.7 ± 28.0 | 57.6 ± 40.0 | 27.6 ± 10.1 |
| *INFB* | 57.7 ± 38.0 | 73.0 ± 35.0 | 66.6 ± 33.5 | 61.5 ± 38.5 | 53.8 ± 19.4 | 79.7 ± 32.3 | 93.3 ± 38.6 | 82.8 ± 28.4 | 65.8 ± 36.3 | 91.6 ± 30.9 | 1002.3 ± 583.7 |
| *INFG* | 6.1 ± 1.8 | 14.2 ± 3.6 | 6.6 ± 1.4 | 7.3 ± 2.2 | 7.2 ± 0.7 | 7.5 ± 2.2 | 5.5 ± 2.1 | 8.6 ± 1.1 | 18.8 ± 1.4 | 14.0 ± 9.0 | 1.76 ± 0.4 |
| *IFNL* | 46.0 ± 37.5 | 52.3 ± 27.2 | 34.9 ± 17.4 | 34.3 ± 24.2 | 30.5 ± 13.8 | 45.9 ± 18.8 | 56.5 ± 20.6 | 51.1 ± 25.4 | 45.6 ± 3.8 | 52.9 ± 24.0 | 675.3 ± 152.1 |
| *TGFB* | 4.0 ± .3 | 2.9 ± 0.4 | 3.7 ± 0.3 | 4.1 ± 0.5 | 3.7 ± 0.1 | 3.9 ± 0.5 | 4.2 ± 0.5 | 3.6 ± 0.5 | 3.0 ± 0.3 | 3.4 ± 0.3 | 1.4 ± 0.1 |
| *TNFa* | 33.2 ± 1.4 | 80.5 ± 12.6 | 44.3 ± 4.2 | 40.9 ± 8.8 | 41.7 ± 4.8 | 49.7 ± 1.5 | 47.4 ± 3.1 | 76.8 ± 7.2 | 88.9 ± 8.2 | 85.6 ± 11.3 | 453.2 ± 73.2 |
| *TLR3* | 544.6 ± 302.6 | 616.5 ± 285.7 | 470.3 ± 242.1 | 503.0 ± 288.7 | 488.0 ± 173.3 | 719.4 ± 279.8 | 794.3 ± 345.3 | 655.7 ± 310.1 | 511.6 ± 235.0 | 494.4 ± 274.9 | 6.1 ± 3.7 |
| *TLR4* | 0.9 ± 0.04 | 0.7 ± 0.07 | 0.8 ± 0.06 | 0.9 ± 0.05 | 0.7 ± 0.05 | 0.9 ± 0.09 | 1.0 ± 0.08 | 0.7 ± 0.1 | 0.5 ± 0.05 | 0.6 ± 0.1 | 0.2 ± 0.02 |
| *TLR7* | 7.8 ± 1.4 | 6.3 ± 0.9 | 5.4 ± 1.0 | 6.5 ± 0.8 | 5.3 ± 0.5 | 8.1 ± 0.8 | 9.3 ± 1.3 | 5.9 ± 0.5 | 5.2 ± 0.7 | 4.9 ± 0.5 | 4.5 ± 1.8 |
| *TLR8* | 43.4 ± 6.8 | 50.6 ± 8.7 | 44.4 ± 6.6 | 42.7 ± 5.2 | 42.7 ± 8.8 | 48.7 ± 9.1 | 47.9 ± 10.6 | 40.8 ± 5.1 | 38.2 ± 6.2 | 37.2 ± 1.9 | 2.1 ± .05 |
| *TLR9* | 2.2 ± 0.3 | 2.3 ± 0.4 | 2.1 ± 0.5 | 2.1 ± 0.4 | 2.1 ± 0.2 | 2.1 ± 0.2 | 2.4 ± 0.6 | 2.3 ± 0.5 | 2.7 ± 0.7 | 3.2 ± 0.5 | 2.3 ± 0.9 |

Mean ± SD
